# Regulation of individual differences in recruitment behaviour within honey bee foraging groups

**DOI:** 10.1101/241679

**Authors:** Ebi Antony George, Axel Brockmann

**Affiliations:** National Centre for Biological Sciences, Tata Institute of Fundamental Research; Bangalore 560056; India

**Keywords:** Social insects, Division of labour, Dance communication, Repeatability of behaviour, Social regulation, Response thresholds

## Abstract

Division of labour is a hallmark of eusocial insect colonies, with different groups of workers engaged in different tasks at the same time. Foraging is a task done by older workers in honey bee colonies. Foragers use the waggle dance behaviour to inform and recruit nest mates to food sources in the environment. The recruitment process incorporates information about the food reward, the colony food stores and the environmental food availability and plays a major role in ensuring efficient exploitation of the food sources available to the colony. However, the role that individual foragers play in driving recruitment is largely unexplored. We observed the dance activity of individual foragers from the same foraging group and showed that there are consistent inter-individual differences within a foraging group leading to a division of labour in the recruitment activity. Next, we studied the effect of changing social interactions on these individual differences. Removing foragers from the group led to an increase in the dance activity of the group of remaining foragers. This was mainly driven by an increase in the dance activity of certain individuals within the foraging groups. In contrast, allowing recruits to join the foraging group had a strong negative effect on the dance activity of all the individual foragers. Our study shows that there is a fine scale division of labour in the recruitment activity within foraging groups and that this is further regulated by changing social interactions. Thus, a complex interplay between individual differences and social interactions drive recruitment activity in honey bees.

## Introduction

Animals must adjust their behaviour to changes in the environment to survive and they do so by using information (signals and cues) from the environment. Individual eusocial insects, living in large colonies, have evolved the capacity to use many different cues and signals to decide which tasks need to be done for the benefit of the colony. The colony’s requirements determine the tasks that need to be done and individual variation in the responsiveness to social cues and signals determine who does the task. Individual differences in responses have a genetic basis and are also experience dependent [1,2].

In honey bee colonies, the basic division of labour is based on age, with older workers engaged in foraging. However, foragers are further divided into scouts and recruits based on how they find the food source, and are also divided into pollen and nectar foragers based on the resource they exploit [3,4]. The regulation of foraging involves foragers as well as groups of workers engaged in other tasks like food receiving and storing [3]. To ensure that the tasks are performed in an efficient manner, there are multiple channels of communication and signalling between these different worker groups [5,6]. As there are no central organizers, every individual in these different groups must independently integrate information from multiple sources to decide and adjust its task. Foraging activity is regulated with respect to the qualities of the food source, colony requirements and the availability of food in the environment. The recruitment behaviour of honey bee foragers, the waggle dance, acts as the main regulatory mechanism of the foraging activity. The waggle dance encodes information about the reward value as well as the spatial location of the food source [7]. This provides the colony with a great degree of flexibility in allocating foragers to multiple food sources especially in heterogenous environments [8–13]. Foragers also integrate cues from receiver bees to incorporate information about the nutritional state of the colony and change their waggle dance activity accordingly [14–17]. A strong negative feedback mechanism, in the form of stop signals from foragers, help in inhibiting the dance activity for overcrowded or hazardous food sources [18–23]. Thus, at the colony level, the waggle dance behaviour of foragers acts as a multi-signal integrator, providing a proxy for both colony and environmental conditions to potential recruits.

There is some evidence that individual foragers differ in their waggle dance activity for the same food source [24,25]. However, there are no detailed studies on individual differences in recruitment behaviour so far and its function in the regulation of recruitment at the colony level so far. In this study, we first observed how consistent individual dance activity is and how much variation there is amongst individual foragers. We then performed manipulative experiments to look at how changes in social cues can affect the dance activity of individual foragers. We manipulated foraging groups by either removing foragers or allowing recruits into the foraging group. Together, our results suggest that there are consistent inter-individual differences in the recruitment behaviour which can be regulated by social cues. Thus, recruitment activity for a food source is regulated by an interplay between individual response thresholds and the social context obtained through interaction with nest mates.

## Materials and methods

### Colony set-up

*Apis mellifera* colonies were obtained from a commercial beekeeper and maintained on the campus of the National Centre for Biological Sciences at Bangalore. All experiments were performed in an outdoor flight enclosure (20m length x 4m width x 4m height) between March 2014 and June 2017. Colonies were placed in a three-frame observation hive and fed with sucrose solution and pollen at two different feeders. A wedge was used near the entrance to direct the foragers to one side of the bottom frame to ensure that this side was used by the foragers as the dance floor [26].

### Experimental procedure

Foragers were trained to a feeder containing 1M sucrose (Fisher Scientific International, Inc., USA) solution for 3-4 days. Foragers regularly visiting the feeder were caught, cold anesthetized and numbered plastic tags were glued to their thorax. One or two days later the experiments were started and conducted either in the morning (0900-1200 hours; 11 experiments) or in the afternoon (1400-1700 hours; 5 experiments), depending on the weather conditions.

#### Consistency experiments

Consistency experiments were done to determine the amount of variation in the dance activity of individual foragers from the same foraging group (i.e., visiting the same feeder). The foraging group size was limited to 7-12 experimental foragers randomly selected from all the foragers that had been marked before. In the experiments, the dance activity of these foragers was observed for 3 hours per day on 3 consecutive days (except in the case of 2 experiments where observations could only be done for 2 consecutive days). During the observation period on each day, the foragers were allowed to feed at an ad libitum feeder filled with 1M, 2M and 1M sucrose solution for the first, second and third hours respectively [24].

Two observers were present at the feeder throughout the experiment. One recorded the number and timing of the foraging trips made by the experimental foragers. The other caught all recruits coming to the feeder to keep the experimental foragers motivated to dance throughout the experiment [7]. These bees were released back to the hive after the observation time. A third observer was present at the hive and video recorded all the dances made by the experimental foragers using a Sony Handycam (HDRCX260/HDRCX240, 1080p recordings at 25/50 fps). In total, 12 different foraging groups from 8 different colonies were tested (table S1).

#### Removal experiments

Removal experiments were done to test whether a change in the composition of the foraging group would affect individual dance activity. These experiments lasted for 6 days and consisted of 2 3-day phases; a Pre and a Post-removal phase with 3-hour observations on each day as described above. The Pre-removal phase was the same as the consistency experiments (in two experiments, this phase consisted of only observations from 2 consecutive days). On the 4^th^ day, shortly before the usual observation time, 2-4 foragers from the foraging group were removed. The remaining foragers were then observed for 3 more days (except in one experiment, were observations could be done only for 2 days after the removal). Twelve different foraging groups from 8 different colonies were tested (in H1 to H8, the most active foragers were removed and in L1 to L4, the least active foragers were removed). The number of foragers to be removed was determined based on the average number of round dance circuits done by the foragers while ensuring that at least 4 foragers were remaining after the removal (table S1).

#### Recruit experiments

In these experiments, the foraging group composition was changed by allowing recruits to join the foraging group. The experiments were done over 2 days with 3-hour observations of the round dance activity of 6-11 experimental foragers on each day. The foragers were allowed to feed at an ad libitum feeder filled with 1M sucrose throughout the observation time. On the first day, all recruits were caught as in previous experiments (Pre-recruits) and on the second day, none of them were caught (Post-recruits). Four different foraging groups from 2 different colonies were tested (table S1).

### Video analysis

Videos were analysed manually using VLC Media Player. The video analysis focussed on the number of round dance circuits done by each experimental forager during each dance. Each round dance circuit consisted of a forager nearly completing a full circular path, with a very short waggling motion of her abdomen towards the end of the path [27].

### Statistical analysis

All statistical analyses were done using R version 3.4.1 [28] (with the RStudio IDE [29]). For each forager, 3 different parameters were calculated for each experimental day: *probability* of dancing (the ratio of the number of dances to the number of trips), *intensity* of dances (the ratio of the total number of dance circuits to the number of dances done) and *circuits/trips* (the ratio of the total number of dance circuits made by each forager to the number of trips made by that forager).

#### Consistency experiments

Repeatability estimates (R) were used to analyse the consistency of individual behaviour in the 3 parameters. Generalized linear mixed effects models (GLMMs) were built with each parameter as the response variable, individual identity as a random effect and a Gaussian error distribution. All individuals from all foraging groups were used in this analysis (n = 117 individuals). Confidence intervals and *p* values associated with the R values were obtained using bootstrapping and permutation respectively (10,000 iterations each). The analysis was done using the rptR package [30].

#### Removal experiments

The analysis of the removal experiments was done at the level of the foraging group and at the level of the individual forager to compare the responses at the two different levels. Only data from those experimental foragers that were remaining after the removal were used in this analysis (76 individuals). Further, at each level, the analysis was done to address two specific questions: 1) which foraging groups/individual foragers showed a change in activity due to the removal and 2) which predictors correlated with this change?

Linear mixed effects models (LMMs) were built to compare the Pre and Post-removal activity of each foraging group/individual forager. The 3 parameters were the response variables, an interaction term between the variable of interest (foraging groups/individual foragers; both categorical variables) and the removal (a categorical variable of 2 levels, Pre and Post) was the predictor and the observation number was a random effect in the models. The parameters were scaled based on individual foragers/foraging groups for the analysis of the foraging groups/individual foragers respectively. Data from each foraging group was analysed separately at the level of the individual forager. LMMs were built using the nlme package [31] and multiple comparisons and *p* value adjustments were done using the glht function in the multcomp package [32].

To identify which predictors correlated with the change in activity shown by foraging groups, model comparisons were done. Sets of generalized linear models (GLMs) were built with the difference in mean activity between Post and Pre-removal condition in the parameters as the response variables (obtained from the previous analysis), different predictors in each model (table S2) and a Gaussian error distribution. A cut-off value of 4 AIC units was used to shortlist models. Only those predictors in these models with a large model averaged effect size and confidence intervals not overlapping zero at the 95% confidence level were considered to be significantly correlated with the difference in activity in the foraging groups. Generalized linear models were built using the base stats package [28] and model selection and averaging was done using the MuMIn package [33] in R.

LMMs were built to test if individual differences in activity after the removal correlated with the Pre-removal ranking (in the 3 parameters and in circuits, dances and trips) amongst the remaining foragers. The response variable was the difference in mean activity between Post and Pre-removal condition of individuals. Separate models were built for each parameter with the same set of six predictors and with the foraging group as a random effect. For each of the 3 parameters, data from only those foraging groups in which at least one forager showed a significant change were used in this analysis. The effect sizes and bootstrapped confidence intervals of each of the predictors were used to determine which predictor had an important effect on the response variable. LMMs were built using the lme4 package [34] in R.

#### Recruit experiments

The analysis of the recruit experiments was done using GLMMs at the level of the foraging group. The data for each of the 3 parameters were zero inflated. So GLMMs were built with each of the parameters as the response variable, presence of recruits (a categorical variable of two levels; Pre and Post) as the predictor, foraging group as a random effect and a fitted Tweedie error distribution. The Tweedie distribution was fitted using the tweedie package [35,36] and GLMMs with a Tweedie distribution were built using the glmmPQL function in the MASS package [37] and the statmod package [38] in R.

## Results

### Consistency experiments

Honey bee foragers showed consistent inter-individual differences in their dance activity, suggesting a ranking of foragers in the dance activity within a foraging group (figure 1). Probability of dancing (R = 0.7014, CI = 0.6148 – 0.7702, *p* = 0.0001), intensity of dances (R = 0.6845, CI = 0.5925 – 0.7552, *p* = 0.0001) and circuits/trips (R = 0.7006, CI = 0.6138 – 0.7692, *p* = 0.0001) all had high repeatability estimates (figure 1 *d*).

**Figure 1:**
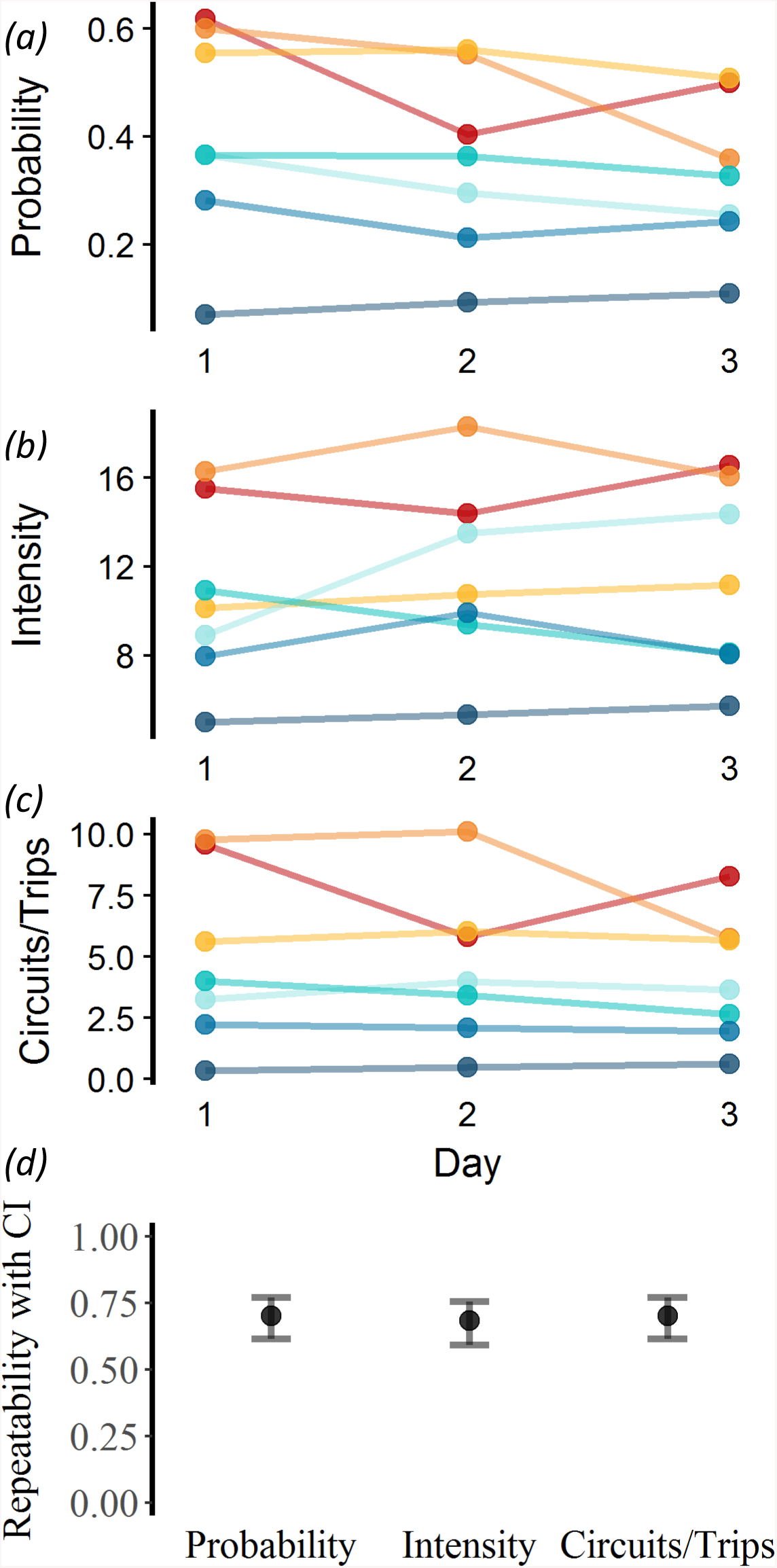
The consistency in activity of 7 foragers from foraging group H3 in all three parameters *(a)* probability of dancing, *(b)* intensity of dances and *(c)* circuits/trips across 3 consecutive days of observation. Each forager is represented by a different colour. *(d)* Repeatability estimates and confidence intervals of all foragers (117 foragers from 12 foraging groups) in all three parameters.

The least and most active foragers strongly differed in their average probability of dancing (0.013 vs 0.706), intensity of dances (2.25 vs 24.02) and circuits/trips (0.06 vs 9.14). Within foraging groups, the mean ratio between the most and least active forager was 3.49 for probability (range = 1.18 - 11.01), 2.24 for intensity (range = 1.31 - 5.77) and 5.41 for circuits/trips (range = 1.22 - 21.15).

### Removal experiments

The removal of foragers from a group led to a change in the average dance activity in 8 out of 12 foraging groups in at least one of the parameters (table 1, figure 2 and figure 3 *a*) irrespective of whether the most active or least active foragers were removed. Five foraging groups (H3, H4, H7, H8 and L2) showed a significant increase in the average scaled probability after the removal. Three foraging groups (H6, L1 and L3) showed a significant decrease in their average scaled intensity after the removal. In circuits/trips, 3 foraging groups (H3, H4 and H7) showed a significant increase while 2 foraging groups (H6 and L1) showed a significant decrease. Four foraging groups (H1, H2, H5 and L4) did not show a significant change in any of the parameters.

**Table 1:** Difference in average scaled activity between Post and Pre-removal conditions for 12 foraging groups in 3 parameters along with single step adjusted *p* values. Foraging groups which show a significant change in at least one parameter as well as the parameter in which they show the change are highlighted in bold.

| Foraging Group | Probability |  | Intensity |  | Circuits/Trips |  |
| --- | --- | --- | --- | --- | --- | --- |
|  | Difference | p Value | Difference | p Value | Difference | p Value |
| H1 | -0.034 | 1.000 | -0.624 | 0.439 | -0.033 | 1.000 |
| H2 | 0.796 | 0.056 | -0.266 | 0.995 | 0.676 | 0.192 |
| **H3** | **1.493** | **<0.001** | -0.131 | 1.000 | **1.203** | **0.004** |
| **H4** | **1.208** | **<0.001** | -0.514 | 0.562 | **0.937** | **0.007** |
| H5 | 0.085 | 1.000 | 0.388 | 0.822 | 0.282 | 0.976 |
| **H6** | -0.449 | 0.501 | **-0.936** | **0.001** | **-0.794** | **0.010** |
| **H7** | **1.280** | **<0.001** | -0.476 | 0.805 | **0.911** | **0.034** |
| **H8** | **1.143** | **0.007** | -0.630 | 0.560 | 0.705 | 0.352 |
| **L1** | -0.372 | 0.917 | **-0.984** | **0.009** | **-0.845** | **0.036** |
| **L2** | **0.691** | **0.039** | -0.142 | 1.000 | 0.520 | 0.293 |
| **L3** | 0.108 | 1.000 | **-1.039** | **0.001** | -0.398 | 0.774 |
| L4 | 0.578 | 0.264 | 0.453 | 0.673 | 0.507 | 0.470 |

**Figure 2:**
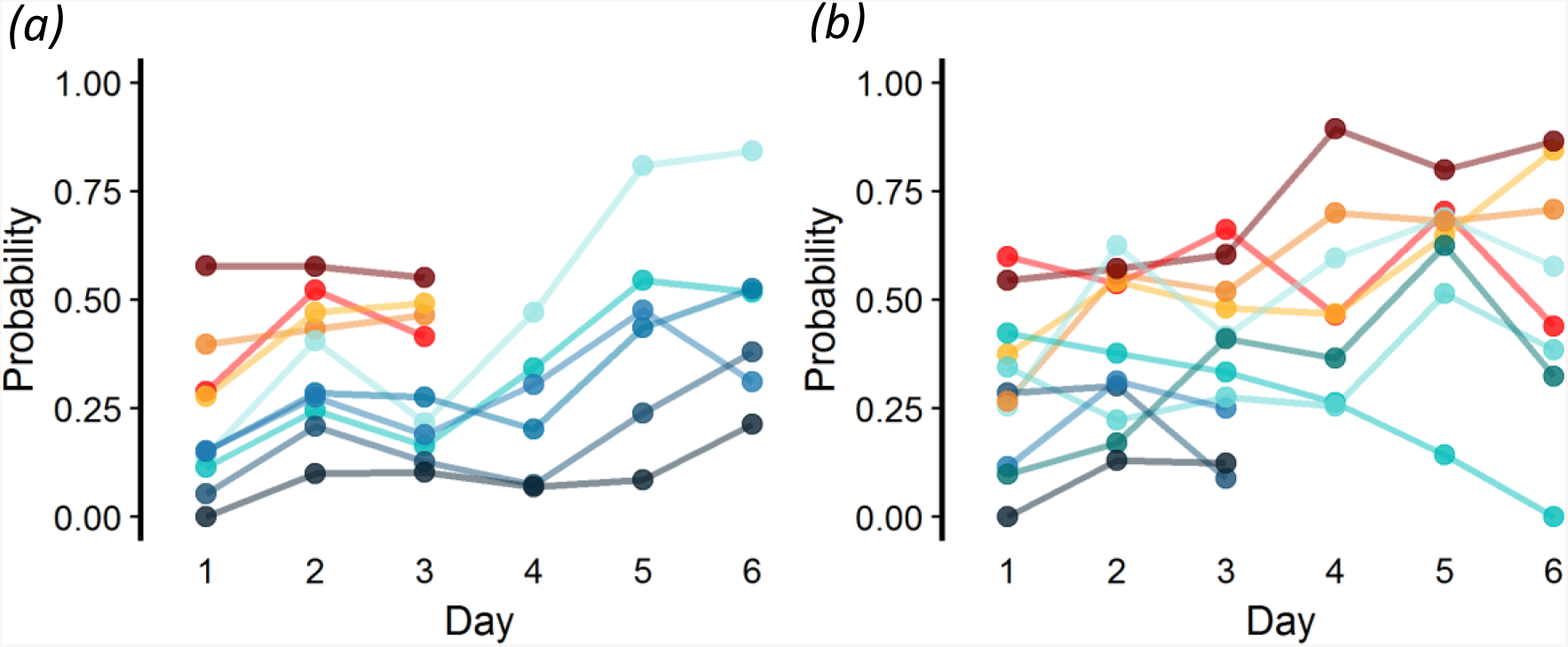
The probability of dancing of foragers from two foraging groups in which removal experiments were done. Each forager within a group is represented by a different colour. *(a)* In foraging group H4, 4 of the most active foragers were removed and 2 out of the remaining 6 foragers showed a significant increase in their probability after the removal. *(b)* In foraging group L2, 4 of the least active foragers were removed and 3 out of the remaining 8 foragers showed a significant increase while one forager showed a significant decrease in their probability after the removal.

**Figure 3:**
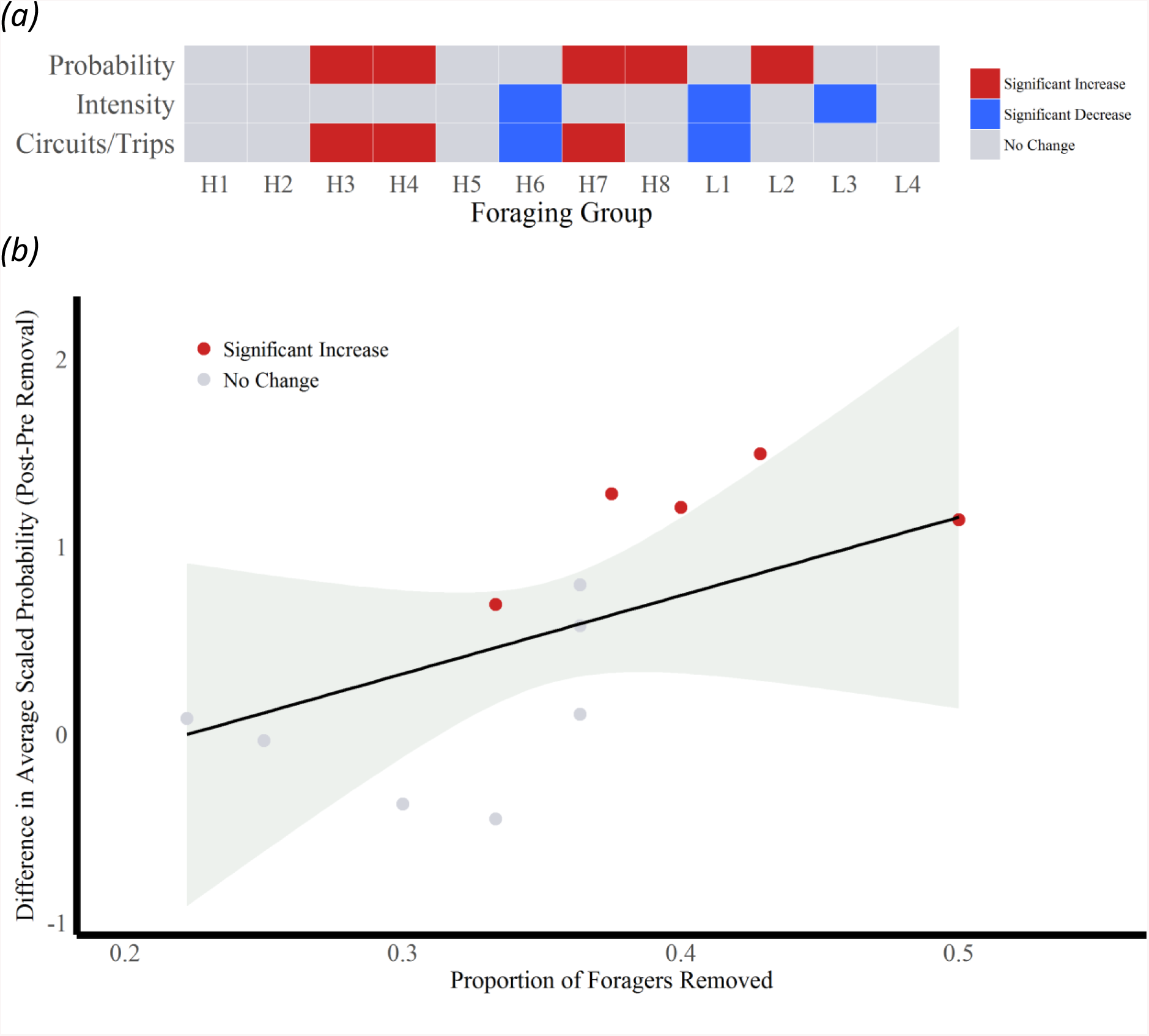
*(a)* Graphical representation of the difference in activity of the 12 foraging groups after the removal in 3 parameters (data in table 1). The foraging groups which show a significant change in activity are highlighted based on whether the change is an increase or decrease in activity. *(b)* The correlation between the difference in probability of dancing due to the removal and the proportion of foragers removed across 12 foraging groups along with the confidence interval. Those foraging groups which show a significant increase are shown as red circles. Increasing proportion of foragers removed led to a greater increase in the probability of dancing of the foraging group after the removal.

In foraging groups, predictors associated with the number of foragers in the group (proportion of foragers removed and foraging group size) most strongly correlated with the change in activity due to the removal. The change in average scaled probability after the removal correlated positively with the number of foragers removed and the proportion of foragers removed (figure 3 *b*), and negatively with the size of the foraging group (table 2). None of the final predictors showed a significant correlation with the change in the average scaled intensity after the removal (table S3). Foraging group size correlated negatively with the difference in the average scaled circuits/trips of the foraging group after the removal (table S4). The proportion of foragers removed had a large positive correlation with this difference but with confidence intervals overlapping zero.

**Table 2:** Important predictors and their effect sizes, confidence intervals, *p* values, relative importance, the number of models the predictor is present in, the AICc, dAIC and cumulative weight of the corresponding model for the difference in probability of foraging groups.

| Predictor | Effect Size | Confidence Interval | p Value | Relative Importance of Predictor | Models Containing Predictor | AICc | dAIC of Model containing Predictor | Cumulative Weight of Model |
| --- | --- | --- | --- | --- | --- | --- | --- | --- |
| Proportion of Foragers Removed | 6.331 | 1.918 – 10.744 | 0.005 | 0.66 | 1 | 23.8 | 0.00 | 0.598 |
| Foraging Group Size | -0.377 | -0.605 – -0.149 | 0.002 | 0.34 | 2 | 25.1 | 1.31 | 0.310 |
| Number of Foragers Removed | 0.692 | 0.199 – 1.185 | 0.006 | 0.34 | 2 | 25.1 | 1.31 | 0.310 |

In 11 out of the 12 foraging groups, there was at least one forager which showed a significant change in its dance activity in at least one of the parameters (table S5 and figure 4 *a*). Even when there was no significant change in dance activity at the group level, there were individual foragers which showed a change in dance activity after the removal (in H1, H2 and L4, but not in H5). Twenty foragers from 10 foraging groups (H1, H2, H3, H4, H7, H8, L1, L2, L3 and L4) showed significant changes in their average scaled probability after the removal (18 foragers showed an increase and 2 showed a decrease). Nine foragers from 5 foraging groups (H1, H6, L1, L2 and L3) showed a significant decrease in their average scaled intensity after the removal. Twelve foragers from 8 foraging groups (H1, H3, H4, H7, L1, L2, L3, L4) showed a change in their average circuits/trips after the removal (9 foragers showed an increase and 3 showed a decrease).

**Figure 4:**
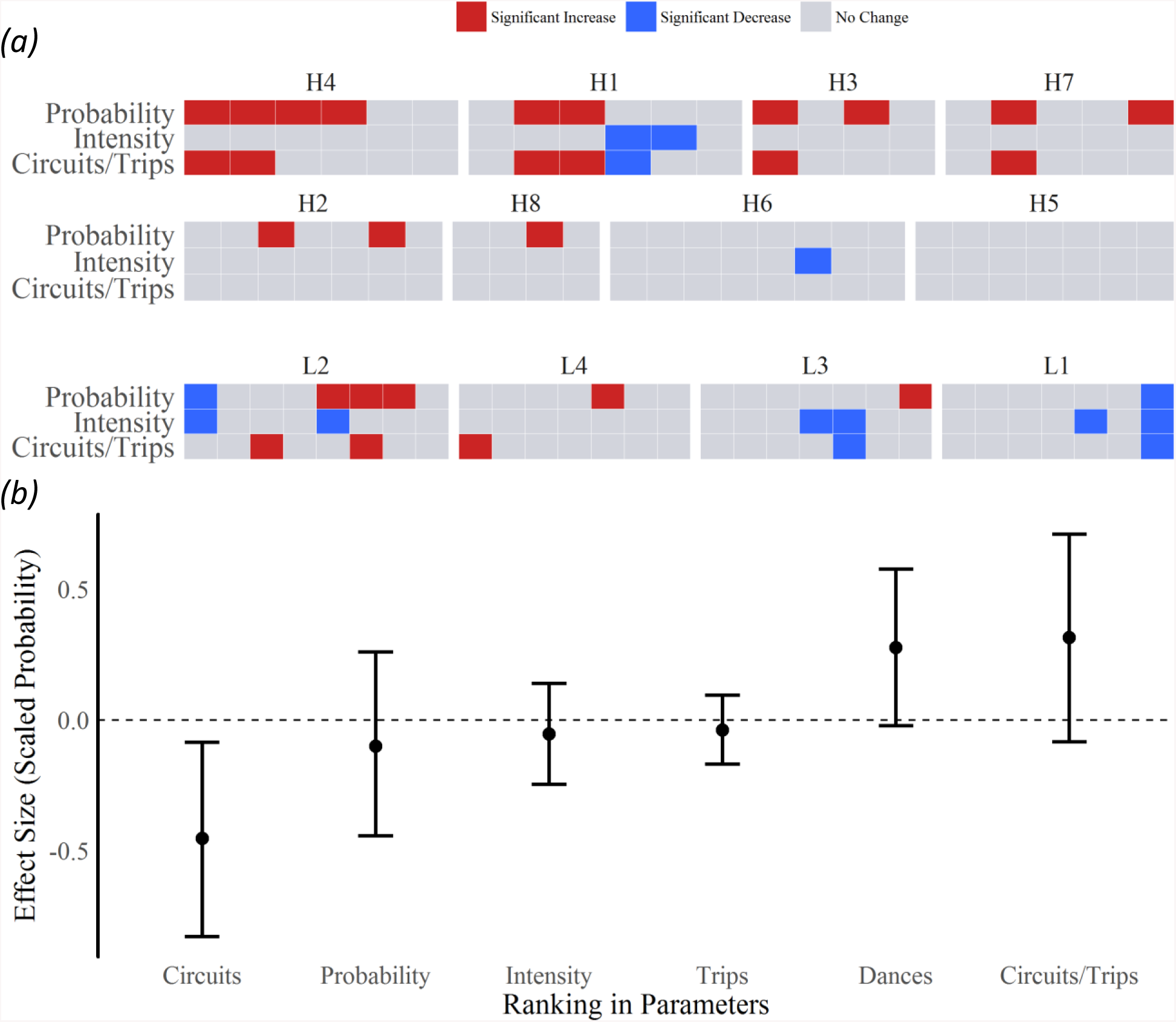
*(a)* Graphical representation of the difference in activity of the 76 remaining foragers from 12 foraging groups after the removal in 3 parameters (data in table S5). The foragers which show a significant change in activity are highlighted based on whether the change is an increase or decrease in activity. *(b)* The effect sizes and confidence intervals of the Pre-removal based ranking in 6 different parameters on the difference in probability of dancing of individual foragers after the removal (data in table 3). Only the ranking in circuits showed a significant correlation with the difference in activity in probability shown by foragers.

The foragers which were highest in their Pre-removal dance activity based rank amongst the remaining foragers showed a greater change in their dance activity due to the removal. The rank of the forager based on circuits in the Pre-removal phase significantly correlated with the difference in dance activity in both the probability and the circuits/trips (table 3 and figure 4 *b*). The ranking in probability, intensity, circuits/trips, dances and trips showed no significant correlation with the difference in either of these 2 parameters. None of the 6 predictors showed a significant correlation with the difference in the intensity of dances due to the removal.

**Table 3:** Effect sizes and bootstrapped confidence intervals of the predictors for the linear mixed effects model with the difference in probability, intensity and circuits/trips of individual foragers as the response variable. Predictors which are correlated significantly are highlighted in bold.

| Predictor<br>(Relative Activity<br>Level in) | Probability |  | Intensity |  | Circuits/Trips |  |
| --- | --- | --- | --- | --- | --- | --- |
|  | Effect<br>Size | Confidence<br>Interval | Effect<br>Size | Confidence<br>Interval | Effect Size | Confidence<br>Interval |
| Probability | -0.101 | -0.444 – 0.259 | -0.064 | -0.586 – 0.432 | -0.038 | -0.444 – 0.422 |
| Intensity | -0.054 | -0.247 – 0.139 | -0.009 | -0.216 – 0.167 | 0.023 | -0.198 – 0.241 |
| Circuits/Trips | 0.316 | -0.085 – 0.710 | 0.127 | -0.356 – 0.597 | 0.210 | -0.219 – 0.706 |
| Trips | -0.040 | -0.169 – 0.094 | -0.065 | -0.179 – 0.049 | -0.052 | -0.191 – 0.067 |
| Dances | 0.276 | -0.023 – 0.578 | 0.060 | -0.308 – 0.413 | 0.229 | -0.167 – 0.566 |
| **Circuits** | **-0.453** | **-0.830 – -0.087** | -0.129 | -0.558 – 0.318 | **-0.465** | **-0.938 - -0.055** |

### Recruit experiments

Allowing recruits to join the foraging group had an immediate strong effect on the dance activity of foragers (figure 5). Most of the experimental foragers (27 out of 32 individuals) completely stopped dancing. The average activity level of the foraging group significantly decreased in probability of dancing (difference = −0.259, *p* < 0.001, power variance index of fitted Tweedie distribution = 1.246), intensity of dances (difference = −8.895, *p* < 0.001, power variance index = 1.03) and circuits/trips (difference = −2.225, *p* < 0.001, power variance index = 1.296). The average activity level of the foraging group when the recruits were present was not significantly different from zero for probability of dancing (mean = 0.0001, *p* = 0.884), intensity of dances (mean = 0.003, *p* = 0.852) and circuits/trips (mean = 0.0004, *p* = 0.92).

**Figure 5:**
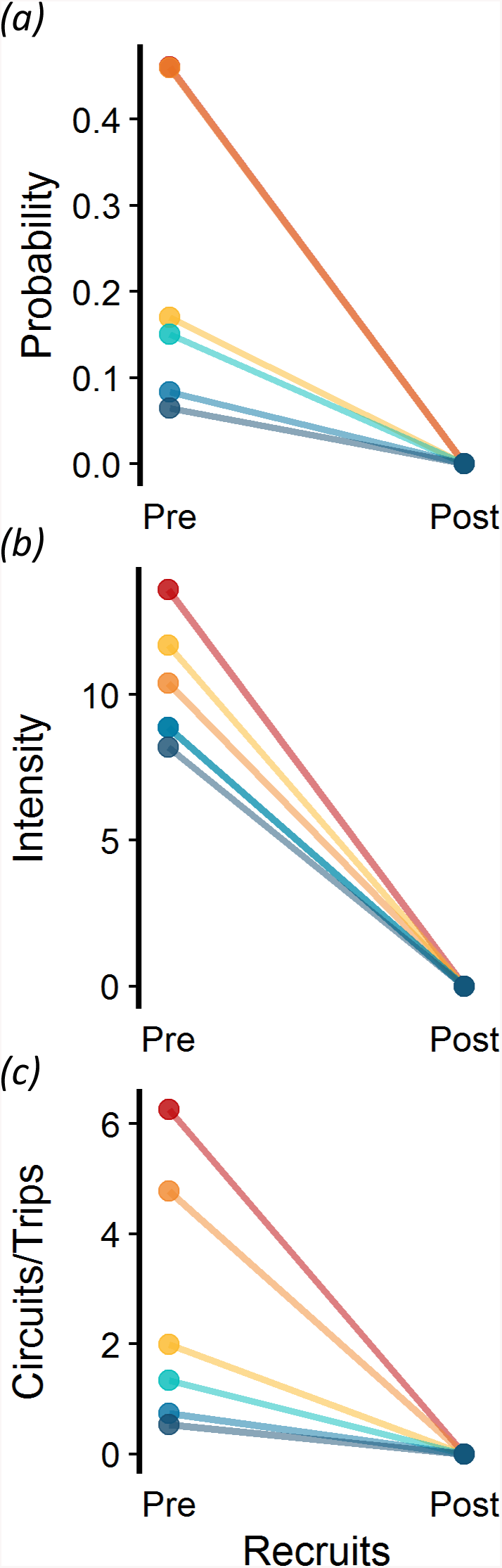
The change in activity of 6 foragers from foraging group A2 after recruits were allowed into the group in all 3 parameters, *(a)* probability of dancing, *(b)* intensity of dances and *(c)* circuits/trips. Allowing recruits into the group led to a stop in the dance activity of all 6 foragers.

## Discussion

The major finding of our study is that honey bee foragers visiting the same food source showed consistent inter-individual differences in their dance activity. These differences were further regulated by the composition of the foraging group. Removal of foragers from the group led to an increase in the probability of dancing of several individuals within a foraging group and the foraging group itself. This increase in probability of the foraging groups positively correlated with the proportion of foragers removed, suggesting that the relative nectar influx into the colony affects the dance activity of foragers. Interestingly, those of the remaining foragers that ranked the highest with respect to the circuits done in the Pre-removal phase showed the greatest increase in their probability after the removal. In contrast, allowing recruits into the foraging group led to a reduction in the dance activity of the whole foraging group and not just specific individuals.

### Consistency experiments

Every honey bee foraging group observed showed a ranking in the dance activity of individual foragers, with some individuals consistently dancing more and others consistently dancing less. Consistent inter-individual differences have been shown for various other invertebrate and vertebrate behaviours before [39–44]. However, apart from a recent study on the repeatability of trapline foraging behaviour [45], there are no other studies on consistent inter-individual differences in honey bee behaviour.

The ranking of individuals with respect to dance activity indicates a fine scale division of labour within foraging groups that has not been identified before. Individual differences in the threshold for evaluating the reward value of the food source might be a mechanism underlying this division of labour [46–50]. There could be different regulatory mechanisms associated with the initiation and the continuation of the dance activity, the combination of which leads to the variation in activity for the same food source. Recent studies have also demonstrated that intra-colonial genetic diversity had a positive effect on dance activity [51].

### Removal experiments

The removal of foragers from the foraging group led to a change in the dance activity at the level of the foraging group and the individual forager. Although only some individuals showed a significant change in dance activity within foraging groups, this led to an overall change in dance activity at the group level. The change in activity at the foraging group level correlated most with the proportion of foragers removed. Foraging groups in which a greater proportion of foragers were removed showed a greater change in dance activity. Within foraging groups, more active individuals showed a greater change in dance activity due to the removal.

Foraging groups showed an increase in their probability or a decrease in their intensity in response to the removal. Increasing the probability rather than the intensity of dances could lead to a more rapid spread of information about the food source to the recruits and hence greater recruitment [52]. This is especially true given the probabilistic nature of recruitment through dancing [53]. The decrease in intensity seen in some foraging groups might indicate that intensity is a noisier representation of changes due to the removal of foragers. Since circuits/trips is related to both probability and intensity, a change in either of these 2 parameters should lead to a change in circuits/trips, although we see this in only 5 out of 8 foraging groups.

The proportion of foragers removed positively correlated with the difference in dance activity in foraging groups. Previous work had shown that the nectar influx has a strong effect on the dance activity at the colony level likely mediated through the interaction between foragers and receivers in the hive [14–17]. The correlation between the proportion of foragers removed and the difference in dance activity suggests that it is the relative and not the absolute decrease in nectar flow that drives the increase in dance activity. Responding to changes in the relative nectar flow might help the colony to be more sensitive to changing environmental food availability and adjust recruitment accordingly [11–13].

The finding that only some individuals from each group responded to the removal of foragers corroborates earlier studies where a similar response was observed in other tasks [54–58]. The higher ranked foragers with respect to circuits done before the removal showed the greatest increase in their dance activity (in probability and circuits/trips) after the removal. It is likely that individual differences in the threshold for responding to changes in receiver interactions might be linked to individual differences in the threshold for evaluating the food reward.

The analysis at the group level and the individual level revealed interesting differences. Even in foraging groups which did not show any change in dance activity due to the removal, there were certain individuals which showed a change in their dance activity. So far, most studies on the honey bee dance communication focussed on responses at the group level [15,59–61]. Our experiments now show that these group level responses do not adequately reflect individual level responses [62]. This serves to further highlight the need for individual level studies on the waggle dance as well as on other behaviours in eusocial insects.

### Recruit experiments

Allowing recruits to join the foraging group led to a drastic reduction in the dance activity in all foraging groups tested. Most experimental foragers in these groups completely stopped their dance activity. In contrast to the removal experiments, the whole foraging group was affected and not just specific individuals. Some of the recruits performed a few waggle dances, but due to a lack of individual specific information this data is not presented.

Most waggle dance experiments performed so far have involved catching and removing recruits to keep the foragers motivated to dance throughout the experiment [7]. Our recruit experiments are more representative of natural foraging conditions wherein recruitment is limited only by the food availability. The results from these experiments suggest that there is an inherent negative feedback within foragers dancing for a food source which is linked to the increasing recruitment to the food source. Limited and decreasing dance activity by an individual is a prominent feature of the waggle dance behaviour for nest sites in honey bee swarms [63,64]. Thus, the dance activity of honey bee foragers for a food source and honey bee scouts for a nest site are more similar than previously appreciated [65].

Overall, our study demonstrates that there is a fine scale division of labour within honey bee foragers visiting the same food source which has not been described before. In addition, the variation in dance activity seen within a foraging group could indicate trade-offs between the time invested in foraging and recruitment, similar to trade-offs between alternative strategies in foraging behaviour in ants [66]. Individual dance activity could be regulated by individual differences in response thresholds for the food reward and the social context along with genetic differences at different levels. This implies that the regulation of foraging at the colony level is a more complicated process than previously known. Further studies need to be done to understand the relationship between response thresholds, individual differences in dance activity and the colony response to changing environmental food availability.

## Authors’ Contributions

Experiments were designed by E.A.G. and A.B. Experiments and data analyses were done by E.A.G. Paper was written by E.A.G. and A.B.

## Acknowledgements

We would like to thank Ravi Boyapati, Abhishek Anand, Hinal Kharva and other student interns for help with the behavioural experiments. We would also like to thank Sruthi Unnikrishnan for providing valuable feedback on the manuscript.

